# Evolutionary dynamics of indels in SARS-CoV-2 spike glycoprotein

**DOI:** 10.1101/2021.07.30.454557

**Authors:** R. Shyama Prasad Rao, Nagib Ahsan, Chunhui Xu, Lingtao Su, Jacob Verburgt, Luca Fornelli, Daisuke Kihara, Dong Xu

## Abstract

SARS-CoV-2, responsible for the current COVID-19 pandemic that claimed over 4.2 million lives, belongs to a class of enveloped viruses that undergo quick evolutionary adjustments under selection pressure. Numerous variants have emerged in SARS-CoV-2 that are currently posing a serious challenge to the global vaccination effort and COVID-19 management. The evolutionary dynamics of this virus are only beginning to be explored. In this work, we have analysed 1.79 million spike glycoprotein sequences of SARS-CoV-2 and found that the virus is fine-tuning the spike with numerous amino acid insertions and deletions (indels). Indels seem to have a selective advantage as the proportions of sequences with indels were steadily increasing over time, currently at over 89%, with similar trends across countries/variants. There were as many as 420 unique indel positions and 447 unique combinations of indels. Despite their high frequency, indels resulted in only minimal alteration, including both gain and loss, of N-glycosylation sites. As indels and point mutations are positively correlated and sequences with indels have significantly more point mutations, they have implications in the context of evolutionary dynamics of the SARS-CoV-2 spike glycoprotein.

## Introduction

Severe acute respiratory syndrome coronavirus 2 (SARS-CoV-2), responsible for the currently ongoing pandemic of coronavirus disease 2019 (COVID-19) (Zhou *et al*., 2020), has infected more than 198 million people and killed over 4.2 million (Anonymous, 2021). Related coronaviruses - SARS-CoV-1 and Middle East respiratory syndrome coronavirus (MERS-CoV) - have also caused pandemics in the recent past.

SARS-CoV-2 belongs to the general class of enveloped viruses (which include influenza and human immunodeficiency viruses among others) that show great plasticity and immune evasiveness due to a protective lipid bilayer and embedded glycoproteins that are heavily N-glycosylated and used as a “glycan shield” (Casalino *et al*., 2020; Cui *et al*., 2009; Watanabe *et al*., 2019). Several genomic features such as point mutations, insertions and deletions (indels), and recombinations impart high diversity within enveloped viruses and profoundly increase their adaptability (Fischer *et al*., 2021). In fact, the envelop glycoproteins of these viruses are particularly variable and found to evolve quickly under selection pressure. As a result, the patterns and drivers of envelop glycoprotein variations in these viruses have been studied extensively (Cui *et al*., 2009; Rao and Wollenweber, 2010b; Rao *et al*., 2015; Zhang *et al*., 2004).

Given the nature of the pandemic, the genomic architecture and its evolutionary dynamics are being keenly explored in coronaviruses (Pavlovic-Lazetic *et al*., 2005; Woo *et al*., 2010) and in SARS-CoV-2 in particular (Badua *et al*., 2021; Li *et al*., 2020b; Lokman *et al*., 2020; Peacock *et al*., 2021). Pachetti *et al*. (2020) have shown the emergence of mutations in SARS-CoV-2 genome and RNA-dependent-RNA polymerase as a mutational hotspot. Mercatelli and Giorgi (2020) have analysed 48635 SARS-CoV-2 complete genomes and found on average 7.23 mutations per sample. Given its importance as a key interactor of angiotensin-converting enzyme 2 (ACE2) for host cell entry and as a target for neutralizing antibodies, spike glycoprotein in the envelop of SARS-CoV-2 is special and therefore its variants are keenly watched (https://www.gisaid.org/hcov19-variants/). D614G was found to be a prevalent spike mutation, however, its precise effect is unclear as it was known to increase infectivity (Korber *et al*., 2020) as well as increase susceptibility to neutralization (Weissman *et al*., 2021). Numerous other mutations in the spike glycoprotein have also been documented (Li *et al*., 2020a).

Studies on variants of SARS-CoV-2 mainly focus on point mutations. This is because there is a massive prevalence of single-nucleotide polymorphisms (SNPs) compared to short indels which only account for 0.8% of mutations (Mercatelli and Giorgi, 2020). A key reason for indels being less common is that they are more deleterious due to frame-shifting compared to SNPs (Choi *et al*., 2012; Lin *et al*., 2017; Mills *et al*., 2006). As SARS-CoV-2 has been shown to accumulate indels (Fischer *et al*., 2021), we are only beginning to explore them and appreciate their myriad roles. For example, Chrisman *et al*. (2021) have looked at indels in the SARS-CoV-2 genome and mapped it to regions of discontinuous transcription breakpoints. Lee *et al*. (2021) have shown a novel indel in nucleocapsid (N) gene leading to negative results for N gene-based RT-PCR that was approved by US/FDA and EU/CE-IVD. Their work also emphasized the genetic variability and rapid evolution of SARS-CoV-2 (Lee *et al*., 2021).

While indels have been explored predominantly at the genomic level (Badua *et al*., 2021; Mercatelli and Giorgi, 2020), they were less emphatically examined at the proteomic level and even less so in the spike glycoprotein. Despite their rarity, indels accelerate protein evolution (Light *et al*., 2013), and could be especially interesting and important in the spike glycoprotein. For example, indels can be beneficial as recurrent deletions in SARS-CoV-2 spike glycoprotein are shown to drive antibody escape and accelerate antigenic evolution (McCarthy *et al*., 2021).

Garry and Gallaher (2021) have explored naturally occurring indels in multiple coronavirus spike proteins and provided evidence against a laboratory origin of SARS-CoV-2. Garry *et al*. (2021) have also shown that the mutations in ‘variants of concern’ (VOC) commonly occur near indels. Despite these revelations, given an accumulating wealth of SARS-CoV-2 genomic data, large-scale patterns of indels in spike glycoproteins have not been fully explored and appreciated.

With this background, we sought to answer some of the open questions: (1) What is the prevalence and pattern of indels in the spike protein? (2) What are the evolutionary dynamics of sequences with indels? (3) Is there any relationship with point mutations? and (4) What is the effect of indels on N-glycosylation sites? We analysed a large set of 1.79 million SARS-CoV-2 spike protein sequences and show that over 50% contained one or more indels. The proportion of sequences with indels has risen sharply, and currently over 76% of unique sequence variants and 89% of total sequences have indels. Indels and point mutations are positively correlated and sequences with indels seem to have more point mutations overall. Further, indels had minimal effect on N-glycosylation sites. We discuss these findings in the context of evolutionary dynamics of the viral protein.

## Materials and Methods

### Spike sequences and metadata

The SARS-CoV-2 spike protein sequences and related metadata were obtained from the GISAID website (https://www.gisaid.org/; accessed on June 03, 2021) (Shu and McCauley, 2017). The spike protein sequences were based on the translation of the genome after alignment to the reference hCoV-19/Wuhan/WIV04/2019 (EPI_ISL_402124) and were in the fasta format. The associated tsv metadata included date of sample collection, location/country of origin, and clade/lineage information of the virus among other details.

### Sequence analyses

There were a total of 1790224 sequences in the database at the time of access. However, as there were numerous quality issues with the data (Maio *et al*., 2020) many sequences were filtered out. For example, there were 465419 sequences containing X (on average of 82.4 X per sequence) that arose from the translation of low-quality regions and/or ambiguous bases in the genome were excluded. Incomplete sequences based on missing N-terminal and/or C-terminal codons were ignored. As our interest was to look for the pattern of short indels (Lin *et al*., 2017), disproportionately short sequences (for example, 3744 sequences were very short - less than 1000 residues in length) that were missing internal parts possibly due to sequencing/annotation issues were ignored. Finally, sequences with incomplete metadata on the date of sample collection were also excluded.

In the final set of 1311545 spike protein sequences, there were a total of 49118 unique sequences based on 100% identity cut-off (Huang *et al*., 2010) that included all possible variants. For each sequence, a pair-wise alignment with the reference (EPI_ISL_402124) was done using Biopython (Cock *et al*., 2009). The BLOSUM62 was used as the substitution matrix, and gap open and extension penalties were set at 11 and 1 respectively (McGinnis and Madden, 2004). Any gap in the query sequence was considered as a deletion and a gap in the reference was considered as an insertion (Choi *et al*., 2012). Finally, a mismatched residue was considered as a point mutation (or substitution).

To see the temporal dynamics, the proportions of sequences with indels were plotted against the date of sample collection (month-wise). Based on the sequence metadata, country and clade/lineage-specific dynamics of indels were also plotted. For each alignment, the number of indels was enumerated and the positions of indels were mapped to the reference sequence. Further, potential N-glycosylation sites were identified based on the regular pattern of tripeptide sequons NXS or NXT where N is asparagine, S is serine, T is threonine, and X is any amino acid residue, and compared with potential N-glycosylation sites found in the reference sequence (Table 1 and S1) (Cui *et al*., 2009; Rao and Wollenweber, 2010a; Zhang *et al*., 2004). Multiple sequence alignments, where required, were done using Clustal Omega (https://www.ebi.ac.uk/Tools/msa/clustalo/). All sequence analysis and data handling, where specifically not mentioned, were performed in Python; and visualization/graphs were created in Microsoft Excel.

**Table 1.**
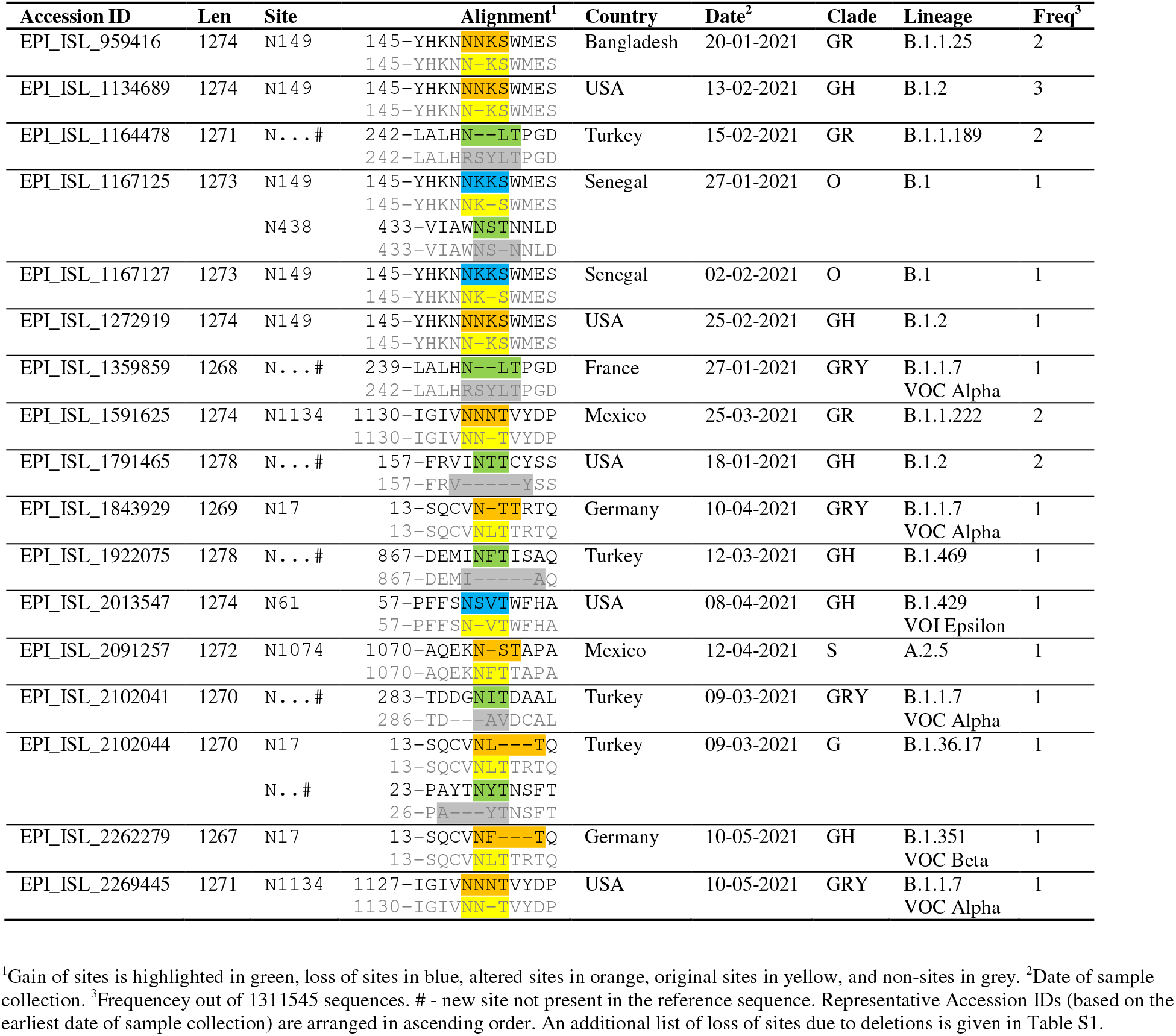
List of N-glycosylation site variants due to indels in SARS-CoV-2 spike protein.

### Structural analyses

The positions of indels and N-glycosylation sites were visualized on the 3D-structure of SARS-CoV-2 spike glycoprotein (PDB ID: 6VXX or 6XR8) using the Visual Molecular Dynamics (VMD) program (https://www.ks.uiuc.edu/Research/vmd/). To see if indels have any preference for sequence/structural/functional features such as surface-exposed regions, solvent accessible surface area (SASA) information was obtained using the DSSP program which calculates an accessibility score (ranged from 0 to 277) from the 3D-structure (http://swift.cmbi.ru.nl/gv/dssp/; Touw *et al*., 2015). Protein disorder was calculated (values ranged from 0 to 0.41) using DISpro (http://scratch.proteomics.ics.uci.edu/), and shorter disorder regions known as molecular recognition features (MoRF) were quantified (values ranged from 0.21 to 0.8) using MoRFCHiBi_Web (https://morf.msl.ubc.ca/index.xhtml) (Malhis *et al*., 2016). Finally, information on different functional domains of SARS-CoV-2 spike glycoprotein was obtained from the literature/UniProt (https://www.uniprot.org/) and residue overlap coefficient was enumerated (Table S2) (Vijaymeena and Kavitha, 2016).

### Statistical analyses

Where required, a one-proportion Z-test was used to check if the observed proportion was significantly different from the expected. A chi-square test for independence was performed to check whether (multiple) sample proportions were significantly different (Agresti, 2007). Correlations between indels and other variables (such as point mutations, accessibility scores, etc.) were measured using more robust Kendall τ coefficient. The significance of correlation coefficient was tested using cor.test(), which is based on t-distribution or approximation. A t-test was used to compare the means of two groups (for example, mutations in sequences with or without indels), and where required, the p-values were corrected for multiple comparison using Benjamini-Hochberg (BH) method (Benjamini and Hochberg, 1995). All statistical tests were done using R.

## Results

### Distribution of sequences with indels

The SARS-CoV-2 spike glycoprotein reference sequence (EPI_ISL_402124) contains 1273 amino acid residues. The distribution of sequences with short indels was plotted based on the number of sequences in each length category, and shown in Fig. 1A as a bar diagram. Overall, the distribution is similar for all sequences (n=1311545, filled bars) and unique sequences (n=49118, open bars). Over 50% of all sequences (inset pie chart) had at least one net indel. It is obvious that sequences with net deletions (50.5%) were far more common compared to sequences with net insertions (0.07%). Further, sequences containing three-residue net deletions were very frequent. A small number of sequences (0.16%) had net deletions of more than three residues. However, these proportions (36.4%, 0.33%, and 1.35%) are significantly different (χ^2^=6991.5, p≈0) if only unique sequences were considered.

**Fig. 1.**
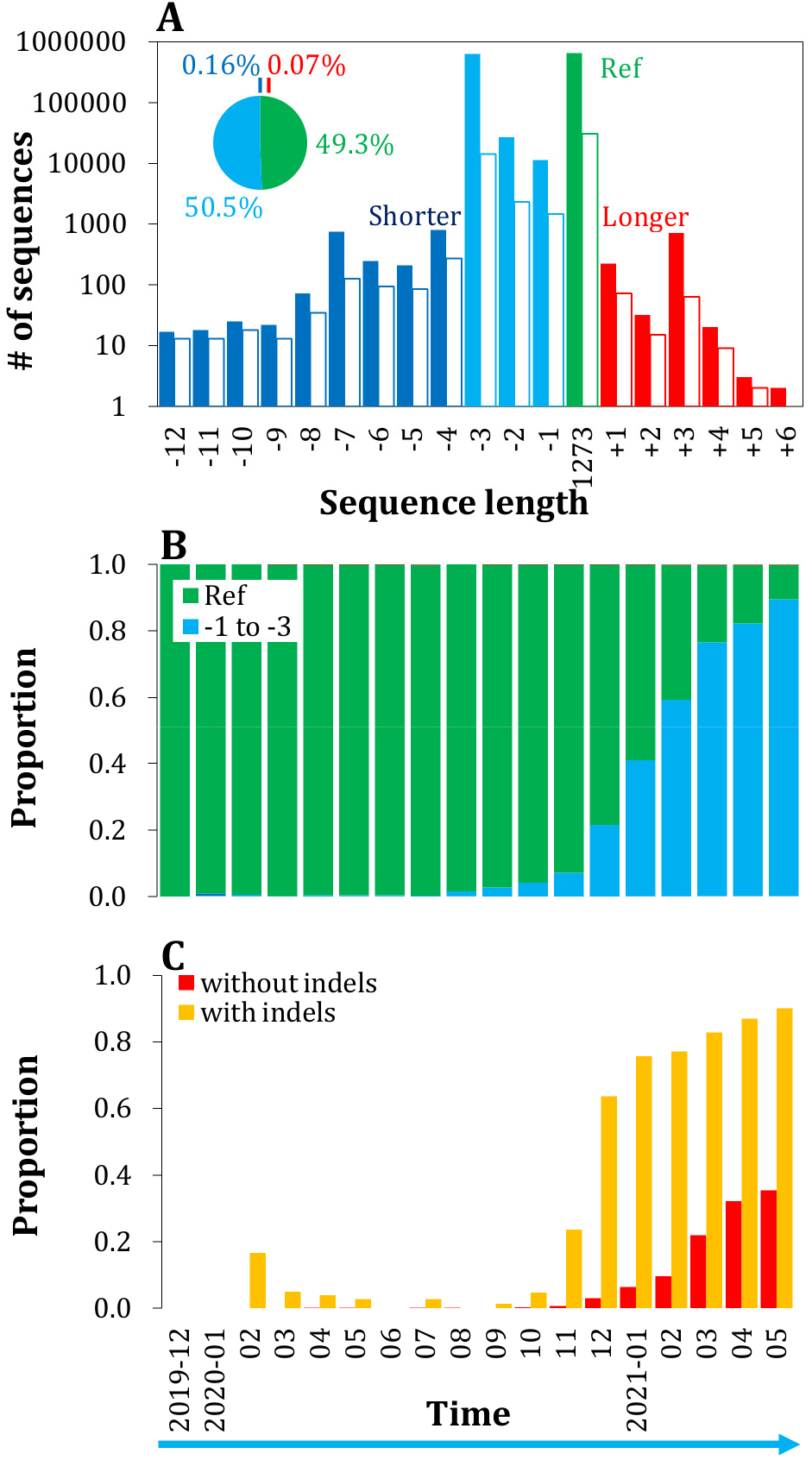
(A) Distribution of sequences with indels in SARS-CoV-2 spike glycoprotein (n=1311545). Over 50% of sequences (inset pie chart) have at least one indel, with sequences containing three net deleted residues being very frequent. The pattern is similar (open bars) even if only unique sequences (n=49118) are considered. (B) Proteins/sequences with indels clearly seem to have a selective advantage as their proportion has risen sharply over time and currently (May 2021) represents 89.3% of all sequences. (C) Month-wise proportions of variants of concern/interest coming from sequences with indels compared to that of without indels.

### Dynamics of sequences with indels

The proportions of sequences with indels over time (month-wise) are given in Fig. 1B, which shows an increasing trend. For example, in August 2020 the proportion of sequences with one to three deletions were about 1.4% which increased to 89.3% in May 2021. The proportions of sequences with insertions, and deletions of more than three residues were also increasing (Fig. S1A). A similar increasing trend is seen even if only unique sequences were considered (Fig. S1B). In May 2021, 76.3% of unique sequences have one to three deletions, and 1.8% of unique sequences with deletions of more than three residues. This increasing trend of sequences with indels holds true across countries (Fig. S1C). Almost all sequence variants currently present in the United Kingdom and South Africa have indels. On the other hand, only a small proportion of sequences from Brazil currently have indels. It should be noted that some patterns were a bit noisy due to small sample sizes (Fig. S2A), for example, in early months and/or country-wise trends. The proportions of sequence contributions from many countries that have reported variants of concern such as India, Brazil, South Africa, and Nigeria were very small (Fig. S2B). In particular, just 0.74% of sequences were from India. However, it becomes 2.7% if only unique sequences were considered (Fig. S2C).

It is interesting to note that a far higher proportion of variants of concern/interest (that have been sharply increasing in the past six months) come from sequences with indels compared to sequences without indels (0.793 vs 0.093, p≈0, two-proportion Z-test). A month-wise trend is shown in Fig. 1C. Overall, 84.1% of unique sequences from VOC/I have one or more indels.

### Incidence of indels along the spike sequence

Figure 2A shows the map of indels along the spike glycoprotein sequence. Figures S3A-C individually show the maps of indels for sequences with net deletions (n=18560), zero net indels (n=92), and net insertions (n=161). While the average number of insertions (2.1, n=161 unique sequences with net insertions) and deletions (2.8, n=18560 unique sequences with net deletions) were small, there were as many as 420 unique indel positions (142 insertion and 358 deletion positions, 80 common positions with insertion or deletion) (Table S2). However, it may be noted that indels were far less common (odds=0.47) in the C-terminal half of the spike protein sequence. Further, indels were frequent only in a few residue-positions. For example, there were just 14 residue-positions (69-70, 140-144, 156-157, 241-243, and 246-247) with 100 or more instances of deletion and only two residue-positions (216 and 217) with 100 or more instances of insertion. Nevertheless, there were as many as 447 unique combinations of indels (ranging from 1 to 15 indels) - for example - insertions at 6, 144, 214-216, etc., and deletions of 69-70_144, 69_70, 144, 156-157, 241-243, etc. Among three-residue deletions, deletion of 69 (amino acid H), 70 (V), and 144 (Y) was the most common combination that has emerged very early in the major lineage B.1.1.7 (VOC Alpha). However, as seen in multiple sequence alignment (Fig. S3D), there is variability in the position of three-residue deletion leading to the emergence of new deletion variants. Fig. 2B shows the multiple sequence alignment of Delta (B.1.617.2) and Kappa (B.1.617.1) variants that contain as many as 17 combinations of deletions (deletion of residues 156 (E) and 157 (F) being the most common). It may be noted that 79% of sequences from Delta and Kappa variants of concern/interest (n=9133, currently represent 0.7% of the total) contain one or more deletions.

**Fig. 2.**
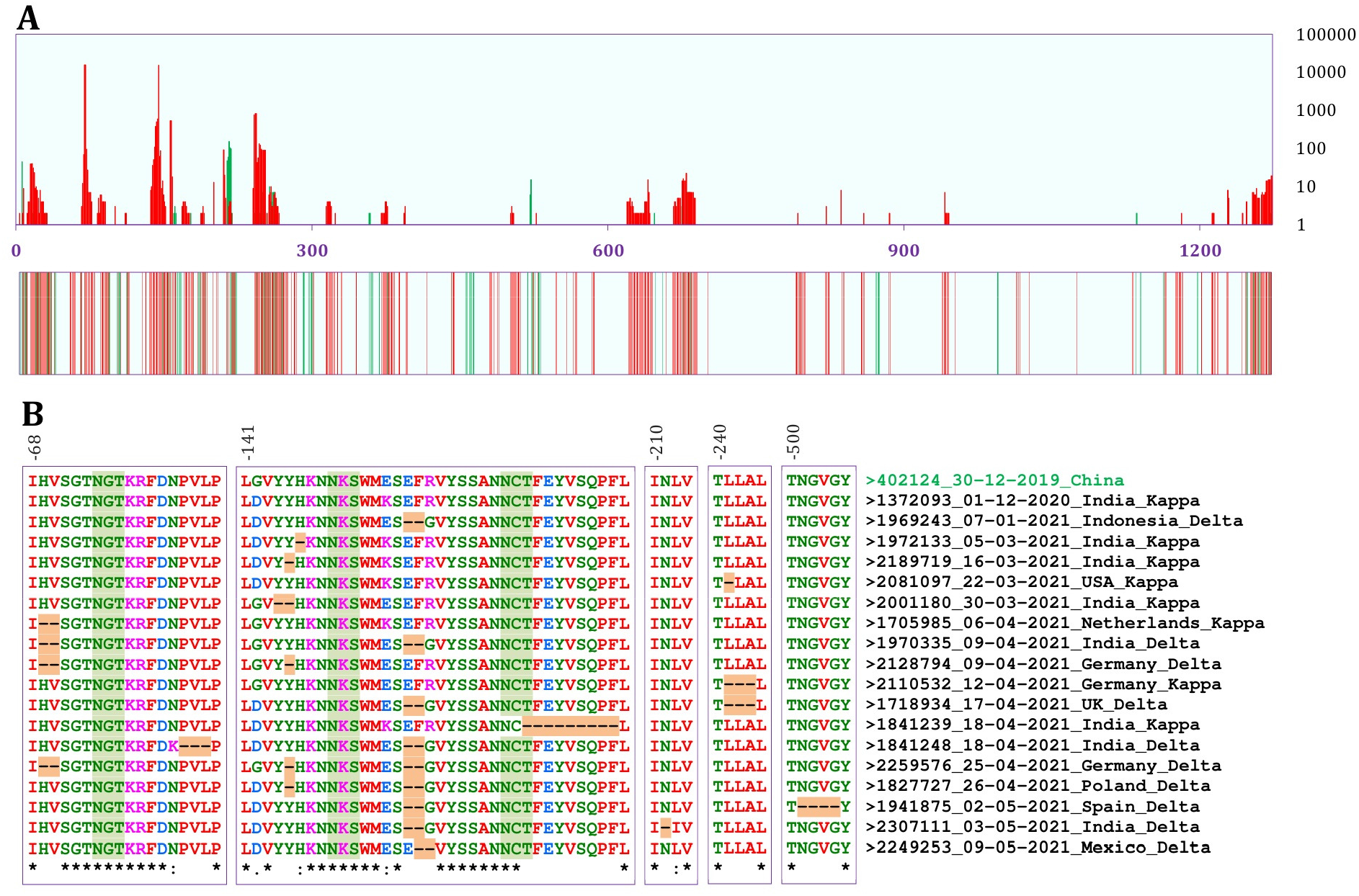
(A) Map of indels (insertion in green and deletion in red) in SARS-CoV-2 spike glycoprotein. Incidence of indels along the sequence. The first panel shows the frequency (scale at right indicates the number of unique sequence variants) and the second panel shows the occurrence of indels. As many as 420 indel positions (142 insertion and 358 deletion positions) are present. Three-residue deletion of 69, 70, and 144 is the most common combination, but there are as many as 447 unique combinations of indels. (B) Multiple sequence alignment (using Clustal Omega, https://www.ebi.ac.uk/Tools/msa/clustalo/) shows 17 combinations of deletions present in Delta (B.1.617.2) and Kappa (B.1.617.1) variants (representative sequences based on the earliest date of sampling). See also Fig. S3.

Based on the correlation, indels showed a low (but significant) preference for surface-exposed regions (τ=0.054 and p=0.027 for insertions, and τ=0.104 and p=1.2E-5 for deletions). Correlation with overall protein disorder was not significant (τ= 0.155 and p=0.56 for insertions, and τ=0.091 and p=0.66 for deletions) possibly because long disordered regions were very few and far apart. Correlation with shorter disordered regions (MoRF) was low but quite significant (τ= 0.122 and p=6.7E-8 for insertions, and τ=0.107 and p=7.1E-7 for deletions). On the 3D-structure of SARS-CoV-2 spike glycoprotein (Fig. S4), indels were prevalent in much of the outer side of the N-terminal domain (NTD). This was reflected in the domain analysis wherein NTD and terminal regions showed a high overlap coefficient (Table S2). Deletions, in particular, were also more frequent at the flanks of receptor-binding domain (RBD), but were far less common in the S2 subunit region and were almost absent at the inner side of the subunits (Fig. S4).

### Indels versus point mutations

The spike sequences with indels had more (over 1.81 times; 7.9±2.1 vs 4.3±2.5, mean ± standard deviation) point mutations compared to sequences without indels. Similar patterns were seen even when different GISAID clades or lineages were taken separately (Fig. 3). However, they were not significant (t-test with BH correction) in some groups when the proportion of sequences with indels was very small or due to a small sample size. Overall, VOC/I had more point mutations compared to non-VOC/I, but in both groups sequences with indels had significantly more point mutations.

**Fig. 3.**
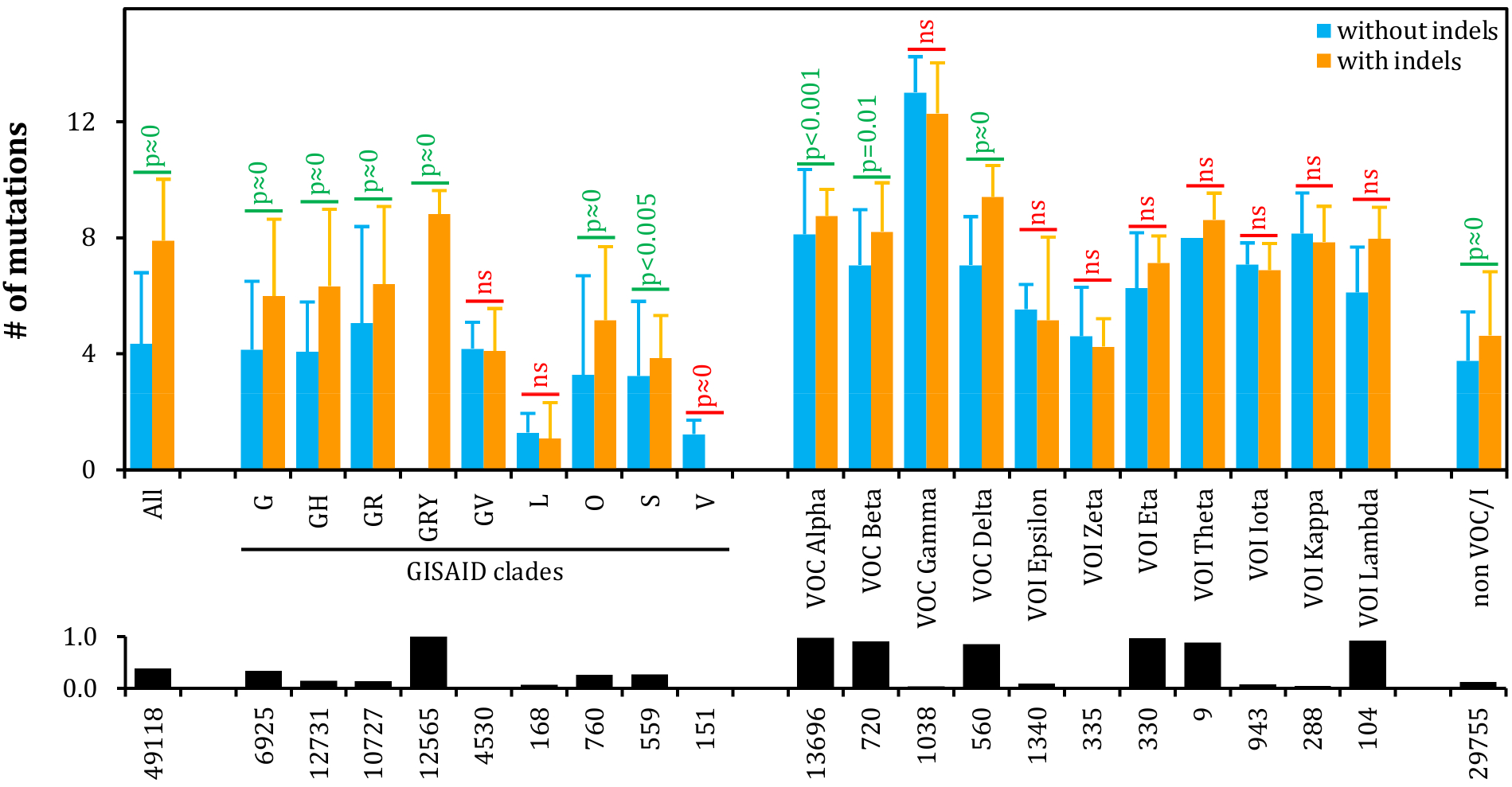
Indels versus point mutations. Bar plot shows the average number of point mutations (mean±SD) in sequences with indels compared to sequences without indels. Point mutations are significantly more (t-test with BH correction) in sequences with indels across clades and lineages of variants of concern/interest. They are not significant (ns, or opposite as in clade V) when sample size and/or proportion of sequences with indels are too small (lower panel of bar plot and sample size).

**Fig. 4.**
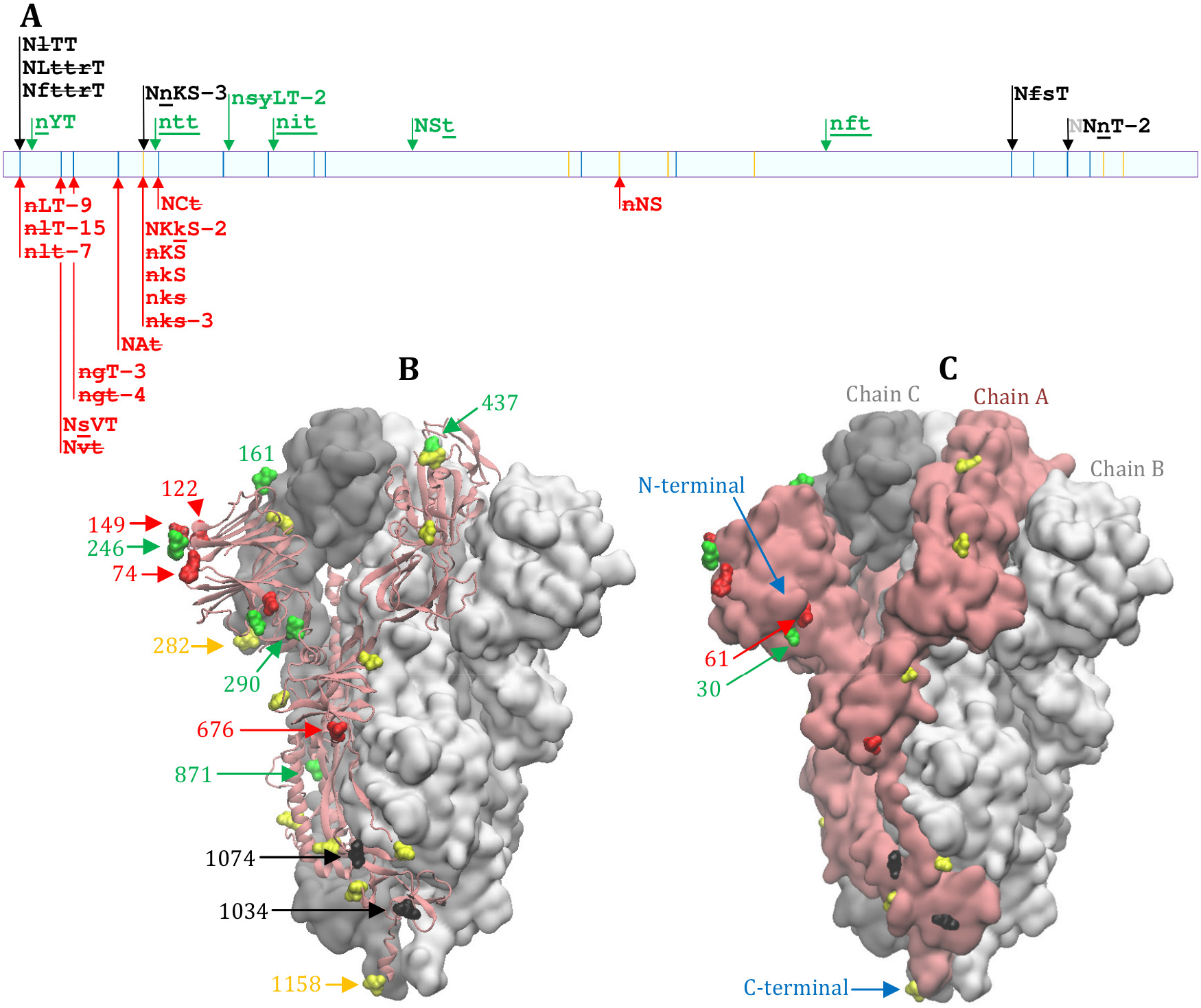
(A) Alteration of N-glycosylation sites due to indels. The panel shows potential N-glycosylation sites (NXS sequons in orange and NXT in blue) along the sequence and the effect of indels (gain of site in green, loss of site in red, and altered site in black). Altered residues (compared to reference sequence) are shown in lower case letters with insertions underscored and deletions struck through. Multiple instances of the same type of change are numbered after hyphen. The gains of sites were more scattered while the losses of sites were mostly at the N-terminal part of the sequence. (B) Cartoon and (C) space-fill structures of SARS-CoV-2 spike glycoprotein showing the positions of N-glycosylation sites. Some key sites are indicated by arrows and numbers. Of the six gains of sites, three sites (at 290, 437, and 871) are completely buried in the 3-D structure.

The distribution of point mutations along the sequence is shown in Fig. S5. Sequences with indels had, apart from D614G, six more very common mutations. There were 96 positions in sequences with indels (n=18813) and 101 in sequences without indels (n=30305, normalized to the number of sequences with indels) that had 100 or more instances of point mutations. Overall, the N-terminal region had longer stretches of residues with more than 100 occurrences of point mutations. There was a small but significant positive correlation (τ=0.224, p=4.0E-25) between the distribution of indels and point mutations along the primary sequence (τ=0.136, p=1.9E-9 for insertions; τ=0.22, p=5.5E-24 for deletions). Many point mutations in VOC Delta were differentially abundant (two-proportion Z-test with BH correction) in sequences with indels compared to sequences without indels (Fig. S5C) and were more common in the N-terminal half where indels are also present (Fig. 2B).

### Effect of indels on N-glycosylation sites

Based on the occurrence of sequons, there are 22 potential N-glycosylation sites (seven NXS and 15 NXT) in the SARS-CoV-2 spike glycoprotein reference sequence (EPI_ISL_402124). Despite indels being present in over 50% of the total 1311545 sequences, there was remarkably minimal effect on N-glycosylation sites due to indels. The list of 67 instances of N-glycosylation sites that have been altered by the indels (in 65 unique sequences) is given in Table 1 (and S1) and their positions are shown in Fig. 3A-C. There were seven instances of gain of sites (Table 1, green) - due to insertions (for example near position 27, `A---YT` to `AYTNYT`), or deletions with substitution (for example near position 246, `RSYLT` to `N--LT`). There were also many more instances of loss of sites (Table 1, blue) - mostly due to deletions (for example at position 17, `VNLT` to `V--T`), but also due to insertions (for example at position 61, `N-VT` to `NSVT`). While the loss of sites occurred mostly at the N-terminal part of the spike, the gain of sites was a bit more scattered. It is important to note that all gains of sites were of NXT types. However, three sites (at around 290, 437, and 871) were completely buried in the 3-D structure (Fig. 3B-C). There were also a few instances of alterations of existing sites due to insertions or deletions (Table 1, orange). It may be noted that these indel-based N-glycosylation site alternations occurred in sequences belonging to many clades/lineages - many of them were of variants of concern (Tables 1 and S1).

## Discussion

There was great interest and urgency to unravel the architecture of the SARS-CoV-2 genome (Zhou *et al*., 2020). Given the severity of the pandemic, there is currently an explosion of viral sequencing bringing along the concerns of data accessibility and ownership (Maxmen, 2021; Noorden, 2021), data integrity/quality (Maio *et al*., 2020), and inequality of sequencing effort/data collection among countries (Colson and Raoult, 2021; Kaur, 2021). Nonetheless, the availability of a vast amount of sequences has allowed the scientific community to track the changes in the SARS-CoV-2 genome as the pandemic is progressing. Numerous studies have shown the emergence and dynamics of new variants, although the main emphasis was on non-synonymous substitutions at the genomic level (Badua *et al*., 2021; Garry *et al*., 2021; Korber *et al*., 2021; Li *et al*., 2020a; Lokman *et al*., 2020; Pachetti *et al*., 2020).

Indel variants were underexplored and unappreciated due to their relative rarity (Badua *et al*., 2021; Mercatelli and Giorgi, 2020; Mills *et al*., 2006), but it is becoming evident that they play key roles in the SARS-CoV-2 genome (Chrisman *et al*., 2021; Lee *et al*., 2021). One reason for indels being even less explored at the proteomic level was because they are primarily found in untranslated regions (Badua *et al*., 2021). In this work, we showed that there is an incursion of short indels at numerous positions in the SARS-CoV-2 spike glycoprotein. Of these, two very common deletions (ΔH69/ΔV70, primarily found in UK variants) were well known and shown to have recurrent emergence and transmission (Kemp *et al*., 2020). While deletions facilitate antibody escape, it was found that BNT162b2 vaccine-elicited sera can still neutralize 69/70 deletion variant (Xie *et al*., 2021). One reason could be concurrent substitutions that offset this effect. For example, D614G substitution, also found in the deletion variant (clade G, prevalent in Europe), was shown to increase SARS-CoV-2 susceptibility to neutralization (Weissman *et al*., 2021). Some independent deletions of five to seven residues were known to occur in and near the furin-like cleavage site (around residue position 681), and it was hypothesized that those deletions might be involved in viral infection (Liu *et al*., 2020). However, at present, the functional implications of numerous other indels are completely unexplored/unknown.

The proportion of proteins/sequences with indels has risen sharply over time. Currently 78.4% of the viral variants have indels; and while Δ69-70 and Δ144 were present in the majority of the variants, there also seems to be an increasing trend for longer indels. Clearly, indels seem to have a selective advantage. Recurrent deletions in the SARS-CoV-2 spike glycoprotein are known to drive antibody escape (McCarthy *et al*., 2021). For example, recurrent deletions (Δ141-144 and Δ146, and Δ243-244) in spike N-terminal domain (NTD) abolished its binding with neutralizing antibody 4A8, and Δ140 caused four-fold reduction in neutralization titre (Harvey *et al*., 2021). The emergence of novel indels leading to variants of concern could be a challenge to vaccines and COVID-19 management (Bian *et al*., 2021; Gupta, 2021; Zhou *et al*., 2021). This could be further exacerbated as sequencing efforts in many (developing) countries were minimal (Colson and Raoult, 2021; Kaur, 2021), but there seem to be disproportionally more variants, for example, in India. The viral diversity/variants may only be fully appreciated if there is better sequencing effort in these courtiers, many of which are reporting the emergence of variants of concern. For example, Resende *et al*. (2021) have found convergent indels in the NTD of spike/SARS-CoV-2 lineages with mutations of concern circulating in Brazil, while Tegally *et al*. (2021) found Δ242-244 in the SARS-CoV-2 variant of concern in South Africa. Recurrent emergence of insertions (between R214 and D215) in the NTD, and their progressive increase in multiple lineages including VOC have been recently documented (Gerdol, 2021). It is important to note that many indel positions are highly variable due to independent/multiple origins of indels (McCarthy *et al*., 2021; Tegally *et al*., 2021) and/or nearby substitutions that affect alignment. Indels have special relevance as they can fine-tune the 3D structure beyond point mutations, and are known to occur in surface-exposed loops (Light *et al*., 2013). As the SARS-CoV-2 will be with us forever (Phillips, 2021), there is a need, equitably across countries, to monitor the dynamics of variants, including indels.

Point mutations (or substitutions) tend to accumulate near the indels (Tian *et al*., 2008; Yang *et al*., 2010). In fact, indels are the driver forces as heterozygosity of indels was proposed as mutagenic to surrounding sequences (Tian *et al*., 2008). As indels are less constrained and have higher structural influence compared to substitutions (Zhang *et al*., 2010), they are frequently under positive selection, for example, in cancer (Yang *et al*., 2010). While the spike mutations in ‘variants of concern’ (VOC) were known to occur near indels (Garry *et al*., 2021), here we showed a large-scale relationship between indels and point mutations. Overall, mutations were more frequent in sequences with indels. This relation holds true even in variants of concern that already have extensive mutations (Wang *et al*., 2021). It is interesting and important to note that sequences with indels have several differentially abundant point mutations in VOC Delta, which is currently posing a global challenge.

Despite numerous instances of indels, there were only a few instances of alterations of N-glycosylation sites in the spike protein. While some existing sites were modified, there were a few more instances of gain and loss of sites. Interestingly, all gains of sites in spike were of NXT type which were known to be preferred by viral glycoproteins (Cui *et al*., 2009; Rao and Wollenweber, 2010a; Rao *et al*., 2011). However, given that some gains of sites were buried in the 3-D structure, they are unlikely to get selected/fixed. Proteins of other enveloped viruses, for example, hemagglutinin (HA) of influenza virus A/H1N1 (since 1918), A/H3N2 (since 1968), and recent A/H5N1 are all accumulating more N-glycosylation sites (Cui *et al*., 2009) and/or modifying the existing sites (Rao and Wollenweber, 2010a; 2010b). It is important to watch the dynamics of N-glycosylation sites in spike as SARS-CoV-2 transforms the vulnerabilities of its glycan shield (Watanabe *et al*., 2020). For instance, the spike protein has 25% glycans by weight which shield approximately 40% of the surface (Grant *et al*., 2020) as against 50% glycans by weight which shield 71-97% in gp120 of HIV-1 countering vaccine development and/or neutralization by antibody (Pancera *et al*., 2014). On the other hand, loss of N-glycosylation sites has selective disadvantage as removal of N331 and N343 drastically reduced infectivity, revealing the importance of glycosylation for viral infectivity (Li *et al*., 2020a). While the SARS-CoV-2 spike utilizes a glycan shield, it also modulates conformational dynamics of the receptor-binding domain by glycosylation. For example, deletion of sites by N165A and N234A mutations reduces spike binding to its receptor ACE2 (Casalino *et al*., 2020).

In conclusion, we show that SARS-CoV-2 is fine-tuning the spike with numerous indels. There seems to have a selective advantage as the proportions of indel variants were steadily increasing over time with similar trends across countries/variants. As many as 420 unique indel positions and 447 unique combinations of indels were present. Indels and point mutations are positively correlated and sequences with indels had significantly more point mutations. Despite their frequency, indels resulted in only minimal alteration of N-glycosylation sites.

## Acknowledgments and Funding

DX acknowledges support by the National Institutes of Health (R35-GM126985). DK acknowledges supports by the National Institutes of Health (R01GM133840, R01GM123055) and the National Science Foundation (DBI2003635, CMMI1825941, and MCB1925643). JV is supported by NIGMS-funded predoctoral fellowship (T32 GM132024). NA acknowledges the initial funding support from the OU VPRP Office for the establishment of the Proteomics Core Facility.

## Data Availability

The SARS-CoV-2 sequences and metadata used in this work are available upon registration, as per the terms of the Database Access Agreement, at GISAID (https://www.gisaid.org/).

## Statement of Ethics

The work is in compliance with ethical standards. No ethical clearance was necessary.

## Author Contributions

RSPR initiated the work and wrote the paper. All authors contributed and were involved in the revision.

## Supplemental information

Supplemental information for this article is available online.

