## Supplemental Information for "Evolutionary dynamics of indels in SARS-CoV-2 spike glycoprotein"

**Table S1.** Loss of N-glycosylation sites due to deletions in SARS-CoV-2 spike protein.

| Accession ID | Len | Site | Alignment[1] | Country | Date[2] | Clade | Lineage | Freq |
|---|---|---|---|---|---|---|---|---|
| EPI_ISL_416884 | 1264 | N74 | 67-A---------KRFD<br>67-AIHVSGTNGTKRFD | Malaysia | 24-01-2020 | L | B | 1 |
| EPI_ISL_434692 | 1265 | N74 | 66-HA--------TKRFD<br>66-HAIHVSGTNGTKRFD | Thailand | 05-01-2020 | L | B | 1 |
| EPI_ISL_459420 | 1268 | N17 | 12-SS-----TTRTQ<br>12-SSQCVNLTTRTQ | United Kingdom | 27-04-2020 | GR | B.1.1.284 | 3 |
| EPI_ISL_513516 | 1269 | N17 | 13-S----LTTRTQ<br>13-SQCVNLTTRTQ | Brazil | 14-04-2020 | GR | B.1.1.33 | 1 |
| EPI_ISL_593557 | 1263 | N17 | 10-LV-------TTRTQ<br>10-LVSSQCVNLTTRTQ | USA | 03-09-2020 | GH | B.1.486 | 1 |
| EPI_ISL_635202 | 1263 | N74 | 67-A---------KRFD<br>67-AIHVSGTNGTKRFD | Slovenia | 07-03-2020 | G | B.1 | 1 |
| EPI_ISL_636459 | 1264 | N74 | 67-A---------KRFD<br>67-AIHVSGTNGTKRFD | Slovenia | 07-03-2020 | G | None | 6 |
| EPI_ISL_793382 | 1266 | N17 | 10-LV-------TTRTQ<br>10-LVSSQCVNLTTRTQ | Denmark | 21-12-2020 | GR | B.1.1.433 | 1 |
| EPI_ISL_878430 | 1266 | N149 | 142-Y-----QSWMES<br>144-YYHKNNKSWMES | United Kingdom | 15-01-2021 | GR | B.1.1.7<br>VOC Alpha | 1 |
| EPI_ISL_901162 | 1268 | N17 | 12-SS-----TTRTQ<br>12-SSQCVNLTTRTQ | Japan | 24-08-2020 | GR | B.1.1.284 | 1 |
| EPI_ISL_952122 | 1265 | N17 | 12-SS-----TTRTQ<br>12-SSQCVNLTTRTQ | United Kingdom | 27-01-2021 | GRY | B.1.1.7<br>VOC Alpha | 9 |
| EPI_ISL_972270 | 1265 | N17 | 11-VS------TTRTQ<br>11-VSSQCVNLTTRTQ | Denmark | 11-01-2021 | G | B.1.258.11 | 1 |
| EPI_ISL_1020465 | 1270 | N74 | 70-VSGS--TKRFD<br>70-VSGTNGTKRFD | Turkey | 26-01-2021 | GR | B.1.1.7<br>VOC Alpha | 1 |
| EPI_ISL_1063795 | 1264 | N17 | 13-S---------Q<br>13-SQCVNLTTRTQ | Spain | 16-02-2021 | GV | B.1.177 | 1 |
| EPI_ISL_1070814 | 1268 | N17 | 13-SQCV--TTRTQ<br>13-SQCVNLTTRTQ | United Kingdom | 03-02-2021 | GRY | B.1.1.7<br>VOC Alpha | 1 |
| EPI_ISL_1088901 | 1271 | N17 | 13-IQCV--TTRTQ<br>13-SQCVNLTTRTQ | USA | 05-02-2021 | GH | B.1.427 | 1 |
| EPI_ISL_1099393 | 1270 | N149 | 145-YH---KSWMES<br>145-YHKNNKSWMES | United Kingdom | 20-02-2021 | GV | B.1.177 | 1 |
| EPI_ISL_1157423 | 1269 | N17 | 12-SS----LTTRTQ<br>12-SSQCVNLTTRTQ | USA | 18-01-2021 | GH | B.1.2 | 1 |
| EPI_ISL_1177349 | 1265 | N17 | 12-SS-----TTRTQ<br>12-SSQCVNLTTRTQ | United Kingdom | 15-02-2021 | GRY | B.1.1.7<br>VOC Alpha | 1 |
| EPI_ISL_1195404 | 1269 | N17 | 13-S----LTTRTQ<br>13-SQCVNLTTRTQ | Germany | 20-11-2020 | GR | B.1.1.37 | 4 |
| EPI_ISL_1209026 | 1263 | N74 | 66-HA---------KRFD<br>66-HAIHVSGTNGTKRFD | Germany | 00-02-2021 | GRY | B.1.1.7<br>VOC Alpha | 1 |
| EPI_ISL_1239453 | 1267 | N17 | 13-S----LTTRTQ<br>13-SQCVNLTTRTQ | USA | 01-03-2021 | GR | B.1.1.7<br>VOC Alpha | 1 |
| EPI_ISL_1239457 | 1267 | N17 | 12-SS----LTTRTQ<br>12-SSQCVNLTTRTQ | USA | 05-03-2021 | GR | B.1.1.7<br>VOC Alpha | 1 |
| EPI_ISL_1295932 | 1272 | N657 | 653-AEHV-NSYECD<br>653-AEHVNNSYECD | Australia | 09-03-2021 | GH | B.1.470 | 1 |
| EPI_ISL_1339912 | 1268 | N17 | 12-SS-----TTRTQ<br>12-SSQCVNLTTRTQ | USA | 10-03-2021 | G | B.1.243 | 1 |
| EPI_ISL_1382125 | 1266 | N17 | 13-S----LTTRTQ<br>13-SQCVNLTTRTQ | USA | 10-03-2021 | GRY | B.1.1.7<br>VOC Alpha | 5 |
| EPI_ISL_1399904 | 1263 | N17 | 13-S-------RTQ<br>13-SQCVNLTTRTQ | Portugal | 01-03-2021 | GRY | B.1.1.7<br>VOC Alpha | 1 |
| EPI_ISL_1472214 | 1269 | N17 | 13-S----LTTRTQ<br>13-SQCVNLTTRTQ | Japan | 26-01-2021 | GR | B.1.1.214 | 11 |
| EPI_ISL_1472219 | 1269 | N17 | 13-S----LTTRTQ<br>13-SQCVNLTTRTQ | Japan | 02-02-2021 | GR | B.1.1.214 | 4 |
| EPI_ISL_1484002 | 1263 | N74 | 65-FH---------TKRFD<br>65-FHAIHVSGTNGTKRFD | United Kingdom | 28-03-2021 | GRY | B.1.1.7<br>VOC Alpha | 6 |
| EPI_ISL_1576799 | 1261 | N17 | 13-S---------Q<br>13-SQCVNLTTRTQ | USA | 26-03-2021 | GRY | B.1.1.7<br>VOC Alpha | 1 |
| EPI_ISL_1585542 | 1268 | N17 | 12-SS-----TTRTQ<br>12-SSQCVNLTTRTQ | Mexico | 18-03-2021 | GR | B.1.1.519 | 1 |
| EPI_ISL_1623710 | 1269 | N149 | 145-YH----NWMES<br>145-YHKNNKSWMES | USA | 07-04-2021 | GH | B.1.526<br>VOI Iota | 1 |
| EPI_ISL_1644014 | 1265 | N17 | 12-SS-----TTRTQ<br>12-SSQCVNLTTRTQ | Germany | 12-04-2021 | GRY | B.1.1.7<br>VOC Alpha | 1 |
| EPI_ISL_1648404 | 1268 | N17 | 12-SS-----TTRTQ<br>12-SSQCVNLTTRTQ | USA | 01-04-2021 | GH | B.1.526<br>VOI Iota | 1 |
| EPI_ISL_1687407 | 1267 | N17 | 11-VS------TTRTQ<br>11-VSSQCVNLTTRTQ | USA | 01-04-2021 | GH | GH_B.1.429<br>VOI Epsilon | 1 |
| EPI_ISL_1728589 | 1262 | N17 | 12-SS--------TQ | Germany | 20-04-2021 | GRY | B.1.1.7 | 4 |

| | | | | | | | | |
|---|---|---|---|---|---|---|---|---|
| | | | 12-SSQCVNLTTRTQ | | | | VOC Alpha | |
| EPI_ISL_1841239 | 1264 | N165 | 161-SSANNC---------L<br>161-SSANNCTFEYVSQPFL | India | 18-04-2021 | G | B.1.617.1<br>VOI Kappa | 1 |
| EPI_ISL_1854607 | 1269 | N17 | 13-SQCV----RTQ<br>13-SQCVNLTTRTQ | Armenia | 30-07-2020 | GR | B.1.1.7<br>VOC Alpha | 1 |
| EPI_ISL_1967942 | 1271 | N17 | 13-SQCV--TTRTQ<br>13-SQCVNLTTRTQ | USA | 25-04-2021 | GH | B.1.526<br>VOI Iota | 1 |
| EPI_ISL_2023476 | 1265 | N149 | 143-V--------WMES<br>143-VYYHKNNKSWMES | USA | 27-04-2021 | GH | B.1.526<br>VOI Iota | 1 |
| EPI_ISL_2138870 | 1265 | N149 | 143-V--------WMES<br>143-VYYHKNNKSWMES | USA | 22-04-2021 | GH | B.1.526<br>VOI Iota | 1 |
| EPI_ISL_2139647 | 1261 | N149 | 141-L------------ES<br>141-LGVYYHKNNKSWMES | USA | 20-04-2021 | GH | B.1.526<br>VOI Iota | 1 |
| EPI_ISL_2145524 | 1271 | N61 | 57-PFFSN--GFHA<br>57-PFFSNVTWFHA | Netherlands | 14-12-2020 | G | B.1.221 | 1 |
| EPI_ISL_2171100 | 1261 | N17 | 13-S---------Q<br>13-SQCVNLTTRTQ | Philippines | 09-03-2021 | GRY | B.1.1.7<br>VOC Alpha | 1 |
| EPI_ISL_2191497 | 1268 | N17 | 12-SS----LTTRTQ<br>12-SSQCVNLTTRTQ | USA | 19-12-2020 | GH | B.1 | 1 |
| EPI_ISL_2256924 | 1269 | N122 | 116-LIVNNA-NVVI<br>118-LIVNNATNVVI | Sweden | 24-03-2021 | GRY | B.1.1.7<br>VOC Alpha | 2 |
| EPI_ISL_2304940 | 1262 | N17 | 12-SS--------TQ<br>12-SSQCVNLTTRTQ | USA | 10-05-2021 | GRY | B.1.1.7<br>VOC Alpha | 2 |

[1]Loss of sites is highlighted in blue and original sites in yellow. [2]Date of sample collection. [3]Frequencey out of 1311545 sequences. Representative Accession IDs (based on the earliest date of sample collection) are arranged in ascending order.

**Table S2.** Distribution of indels in different functional domains of the SARS-CoV-2 spike glycoprotein (Table S2.xlsx).

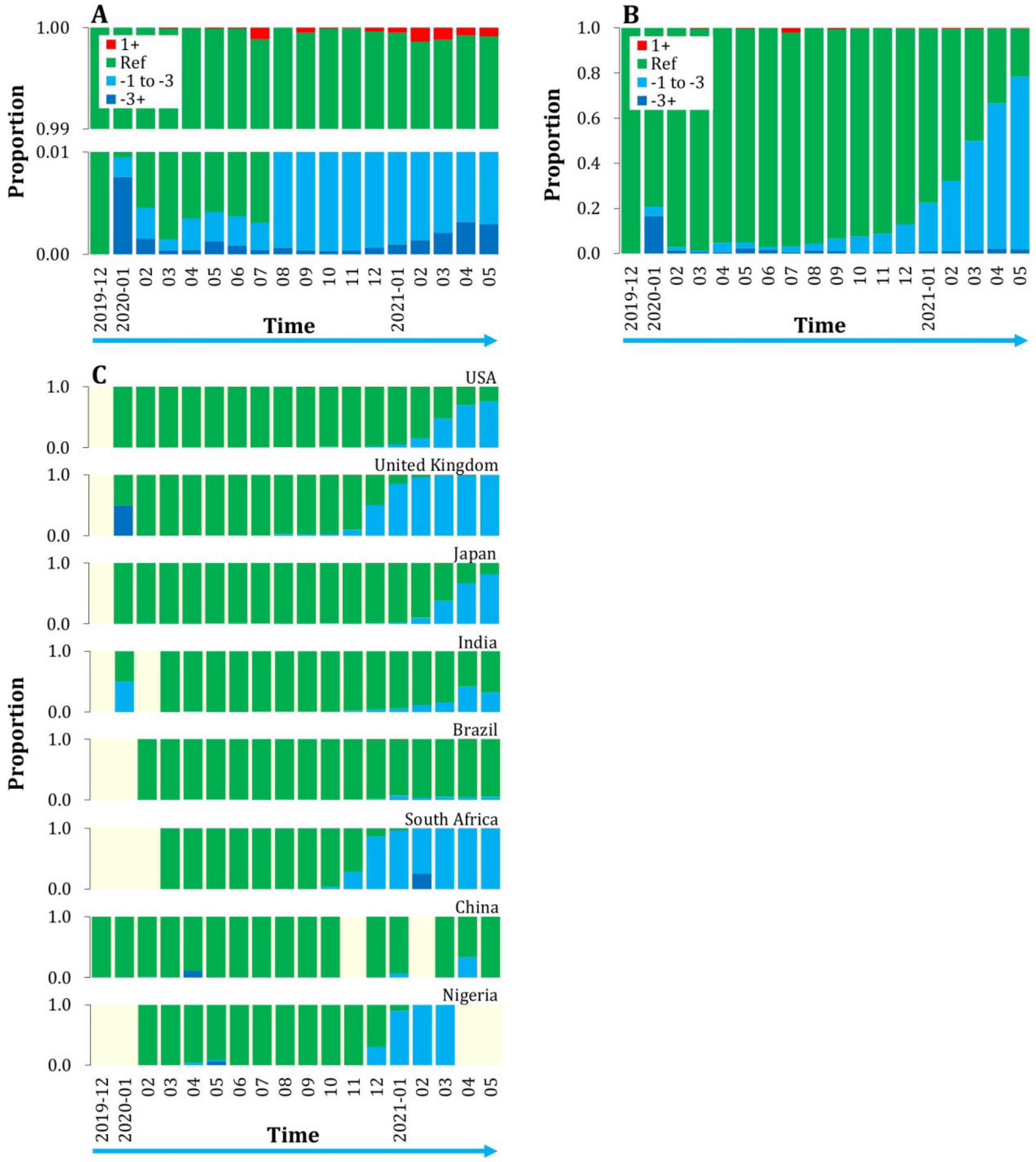


**Fig. S1.** (A) Month-wise proportions of sequences with insertions (red, upper panel) and deletions of more than three amino acids (dark blue, lower panel) also seem to be increasing. However, trends are noisier due to the low number of sequences (see Fig. S2A). (B) Month-wise proportion of sequences with deletions (blue) is clearly increasing even if only unique sequences (n=49118) are considered. (C) Month-wise proportions of sequences with deletions (blue) are increasing across countries including the United Kingdom, India, South Africa, and Nigeria that all have reported the rise of new variants of concern.

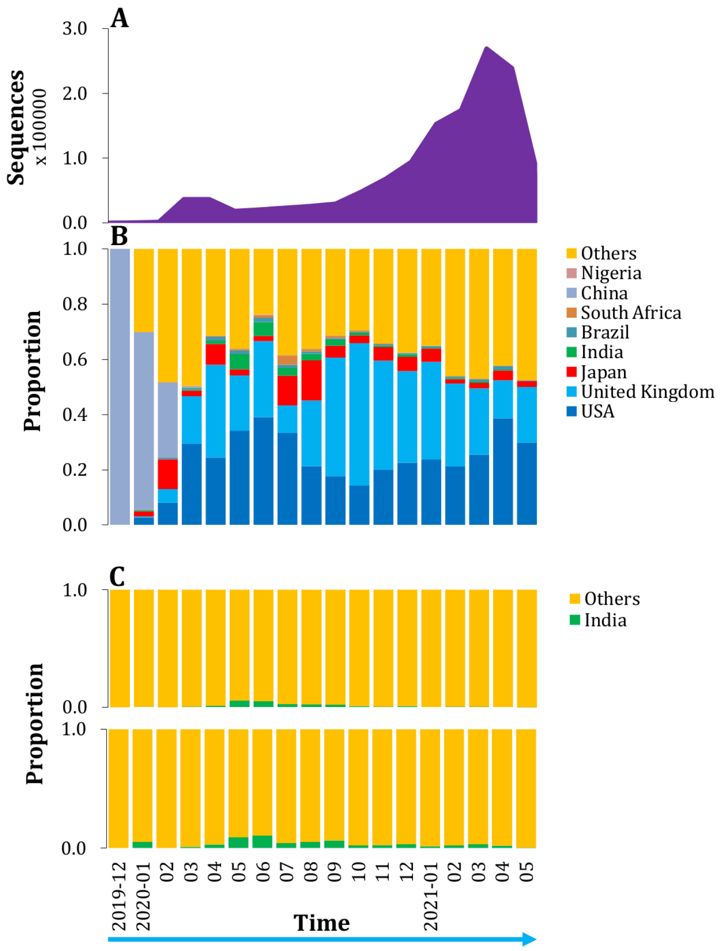


**Fig. S2.** (A) Month-wise number of sequences. (B) Country-wise proportion of sequences. The USA and United Kingdom together account for over 53% of sequences. Others, mainly European countries such as Germany, Denmark, and Sweden contributed large proportions of sequences. Despite the raging pandemic and need for research, India's contribution was paltry. (C) Just 0.73% of sequences were from India which peaked in May/June 2020 (upper panel) and recent monthly proportions of sequences are hardly noticeable. However, India seems to have a disproportionately higher percentage (2.7%) of unique sequences (lower panel), indicating that there could be many more variants given enough sequencing effort.

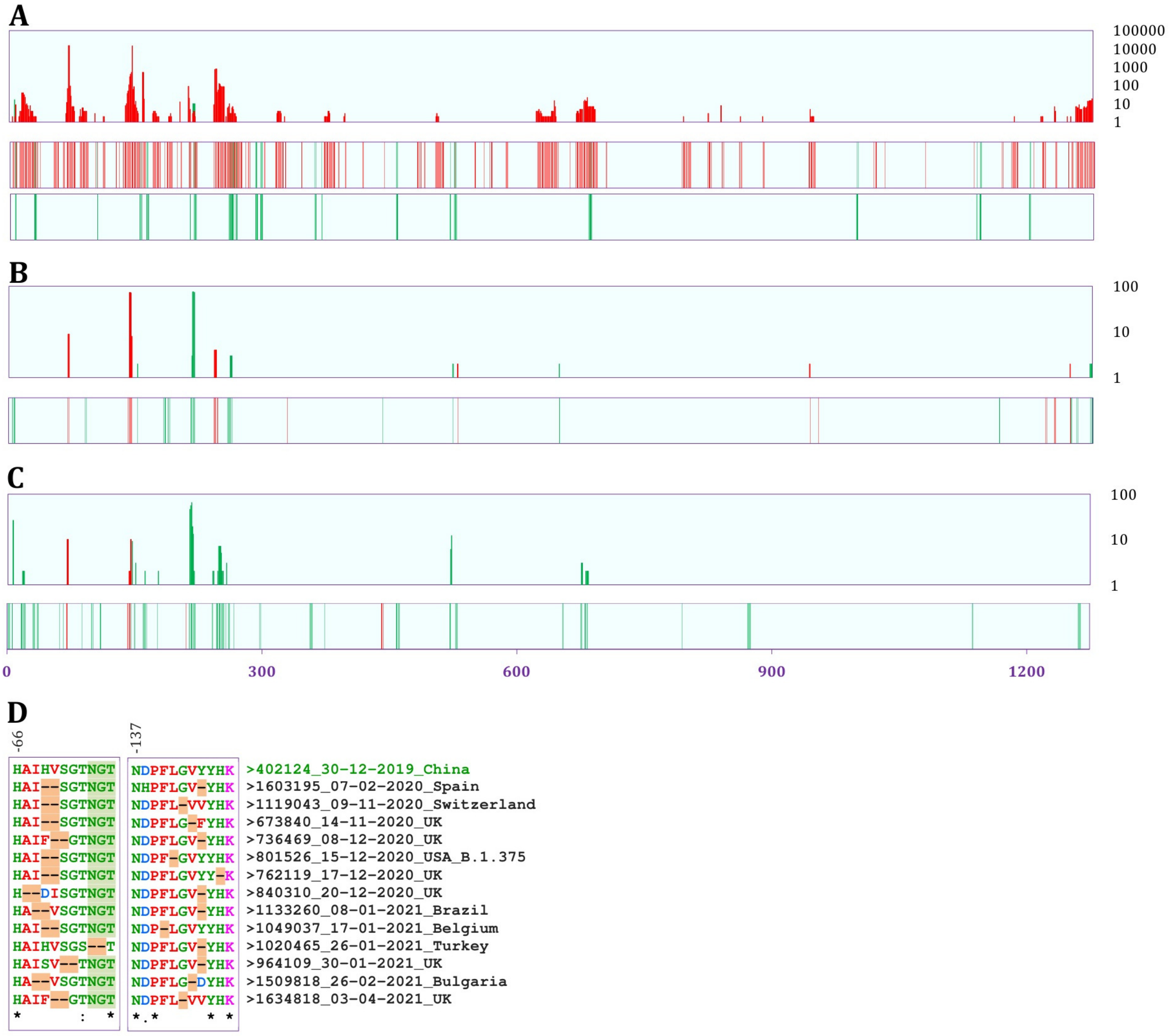


**Fig. S3.** Map of indels (insertion in green and deletion in red) in SARS-CoV-2 spike glycoprotein. Incidence of indels along the length in sequences with (A) net deletions (n=18560 unique variants, which represent 50.7% of total 1311545 sequences), (B) zero net indels (n=92), and (C) net insertions (n=161). The first panels show the frequency (scales at right indicate the number of unique sequence variants) and the second panels show the occurrence of indels. The third panel in (A) shows the insertions which are masked in the second panel. As many as 375 indel positions (48 insertion and 350 deletion positions) are present in (A), 55 (35 and 20) in (B), and 102 (92 and 10) in (C). (D) Among three-residue deletions, deletion of 69, 70, and 144 is very common, but sequences have variability in deletion positions as shown in the multiple sequence alignment (using Clustal Omega, https://www.ebi.ac.uk/Tools/msa/clustalo/) of a few representative variants (based on the earliest date of sampling). Common three-residue deletion variant emerged in 07-02-2020 (lineage B.1.1.7, VOC Alpha), but several new indel variants (mostly in the same lineage) have emerged in the past six months.

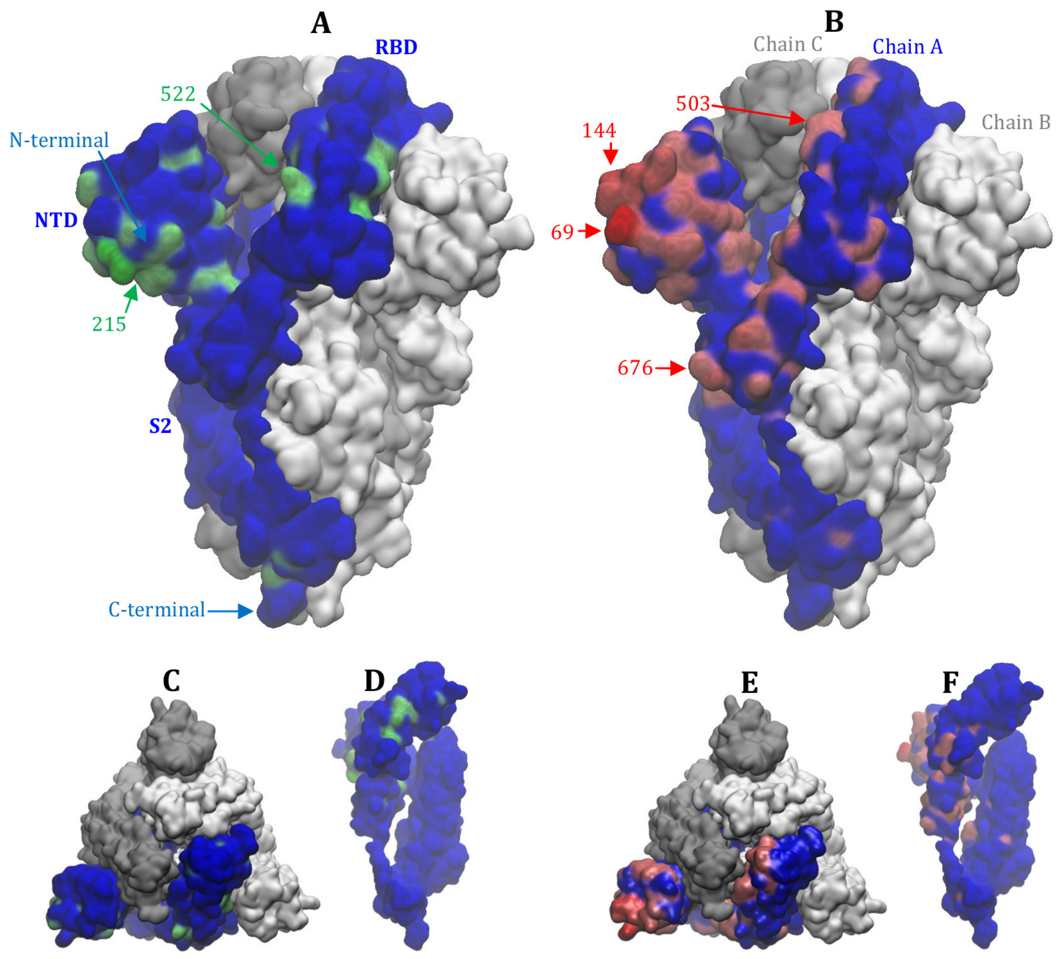


**Fig. S4.** Locations of indels in the 3D-structure of SARS-CoV-2 spike glycoprotein. (A, C, and D) Insertion positions are shown as green and (B, E, and F) deletion positions are shown as red. Indels were prevalent in much of the outer side of the N-terminal domain (NTD). Deletions, in particular, were also more frequent at the flanks of the receptor-binding domain (RBD), but were far less common in the S2 subunit region and were almost absent at the inner side of the subunits. C and E show top views, and D and F show the inner side views of the subunit. A few prominent indel residue positions are shown by arrows and numbers.

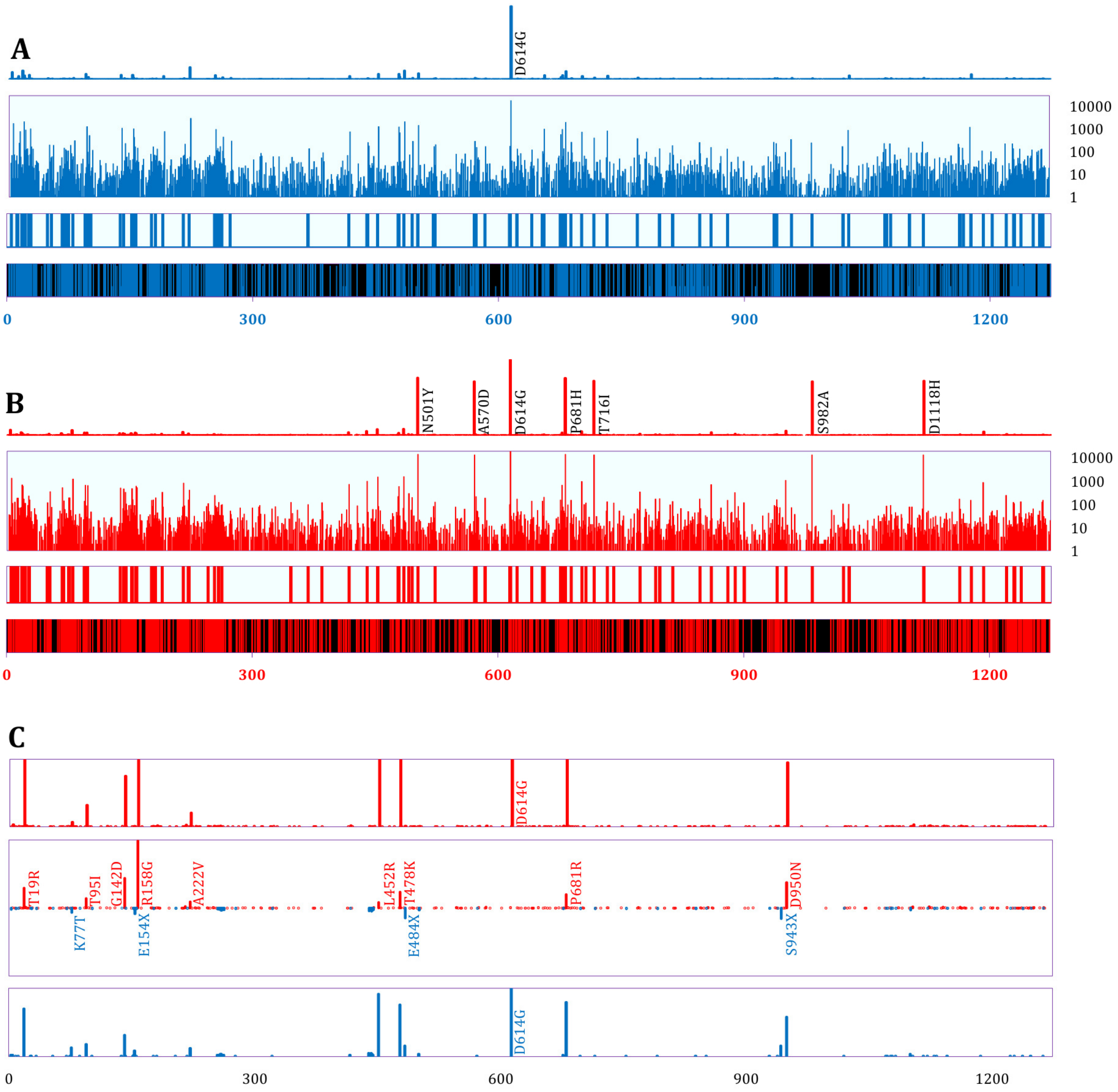


**Fig. S5.** Mutation profile of the SARS-CoV-2 spike glycoprotein. The distribution of point mutations along the length in (A) sequences without indels (n=30305, normalized to the number of sequences with indels) and (B) sequences with indels (n=18813). The top panels show relative mutation frequency (D614G proportion is 0.965 in A and 0.992 in B), and the second panels in log scale highlight the low abundant mutations. Apart from D614G, six more very common mutations present in sequences with indels are mostly coming from VOC Alpha, which is a major lineage. The third panels show residue-positions - 101 in sequences without indels and 96 in sequences with indels - that had 100 or more instances of point mutations. Overall, the N-terminal region had more frequent and longer stretches of residues with more than 100 occurrences of point mutations. The forth panels show regions (black) with 10 or fewer instances of mutations and are more evident between 900 and 1050. (C) Point mutation profile in VOC Delta. The mid panel shows many differentially abundant mutations ($p < 0.05$, two-proportion Z-test with BH correction) between sequences with indels (the top panel) and sequences without indels (the bottom panel), and are more common in the N-terminal half where indels are present.
